# Threshold assessment, categorical perception, and the evolution of reliable signaling

**DOI:** 10.1101/2020.05.30.125518

**Authors:** James H. Peniston, Patrick A. Green, Matthew N. Zipple, Stephen Nowicki

## Abstract

Animals often use assessment signals to communicate information about their quality to a variety of receivers, including potential mates, competitors, and predators. But what maintains reliable signaling and prevents signalers from signaling a better quality than they actually have? Previous work has shown that reliable signaling can be maintained if signalers pay fitness costs for signaling at different intensities and these costs are greater for lower quality individuals than higher quality ones. Models supporting this idea typically assume that continuous variation in signal intensity is perceived as such by receivers. In many organisms, however, receivers have threshold responses to signals, in which they respond to a signal if it is above a threshold value and do not respond if the signal is below the threshold value. Here, we use both analytical and individual-based models to investigate how such threshold responses affect the reliability of assessment signals. We show that reliable signaling systems can break down when receivers have an invariant threshold response, but reliable signaling can be rescued if there is variation among receivers in the location of their threshold boundary. Our models provide an important step towards understanding signal evolution when receivers have threshold responses to continuous signal variation.

## Introduction

In contexts ranging from mate choice to aggression, animals use signals to assess each other (Maynard Smith and Harper 2003; Searcy and Nowicki 2005; Seyfarth et al. 2010). These signals can include any behavioral or morphological trait of one individual (the “signaler”) that has evolved to convey information to another individual (the “receiver”), such as the song of a bird, the sex pheromone of a moth, or the aposematic coloration of a poison frog (Bradbury and Vehrencamp 2011). The information conveyed in signals often regards the quality of the signaler, such as the signaler’s size (e.g., call frequency in frogs; Ryan 2001) or physiological condition (e.g., plumage brightness in some birds; Lindström and Lundström 2000). A central question for researchers studying animal signaling systems is, what maintains the reliability of assessment signals (Searcy and Nowicki 2005)? That is, if it benefits signalers to produce a signal indicating a better quality than they actually have, why do signalers produce a signal that reliably (“honestly”) communicates quality?

In a foundational verbal model, Zahavi (1975) suggested that assessment signals could be reliable indicators of quality if signalers pay costs for expressing signals. This idea was termed the “handicap principle” in that signalers of the highest quality could afford to pay greater handicaps to produce a signal than could signalers of lower quality. Although reliable signaling was shown not to evolve when Zahavi’s original assumptions were made explicit in genetic models (Maynard Smith 1976; Kirkpatrick 1986), Grafen (1990*a*, 1990*b*) later showed, using both population genetic and game theory models, that the handicap principle can lead to the evolution of reliable signaling if two conditions are met. First, the costs of signaling at a given intensity (e.g., producing a large or colorful signal) must be greater for signalers of lower quality compared to signalers of higher quality. Second, receivers should be able to assess continuous variation in signal intensities and thereby gauge signaler quality; that is, as signal intensity increases continuously, so does the benefit the signaler gains from receivers. (Note that Penn and Számadó (2020) suggested that Grafen’s models are substantively different from Zahavi’s original conceptualization. Here, we refer to them both as the “handicap principle” for consistency with the literature). Johnstone (1997) later developed graphical versions of Grafen’s (1990*a*, 1990*b*) mathematical models, illustrating how the optimal level of signaling—the equilibrium point at which the net benefits are greatest—will be different for lower *versus* higher quality signalers. This variation in equilibrium points leads to reliable signaling (Johnstone 1997).

The cost-based reliable signaling theory developed by Zahavi (1975), Grafen (1990*a*, 1990*b*), and Johnstone (1997) has played a crucial role in our understanding of the evolution of animal signals (Maynard Smith and Harper 2003; Searcy and Nowicki 2005). Other studies have recognized, however, that some assumptions of these models do not necessarily reflect the reality of animal signaling systems. For example, the initial models of Grafen (1990*a*, 1990*b*) did not incorporate the concept of perceptual error—when a signaler’s true signal value is not perceived as such by a receiver. In a subsequent series of models, Grafen and Johnstone showed that adding perceptual error into models of signal evolution makes otherwise continuous signaling systems more discrete, such that fewer equilibrium signaling values exist (Johnstone and Grafen 1992a; Grafen and Johnstone 1993; Johnstone 1994). Reliable signaling can be maintained in these models as long as signalers with greater signal intensity are more likely to be perceived as having higher-intensity signals (Johnstone and Grafen 1992a). Furthermore, a continuum of equilibrium signaling values reemerges if perceptual errors are more common at higher signal intensities or if signaling costs increase more rapidly at higher signal intensities (Johnstone 1994). Other researchers have found results similar to those of Grafen and Johnstone when incorporating other aspects of biological realism into models of signal evolution: when the assumptions of the classic handicap models are relaxed, signaling systems are altered, but reliability can be maintained (e.g., Lachmann and Bergstrom 1998; Proulx 2001).

The aforementioned studies have advanced our understanding of signal evolution, but they have all maintained the assumption that receivers can assess continuous variation in signal intensities and thereby gauge signaler quality; that is, as signal intensity increases continuously, so does the benefit the signaler gains from receivers. However, some organisms have threshold responses to assessment signals in which signal receivers respond in a binary manner to continuous variation in signal intensity (e.g., Masataka 1983; Zuk et al. 1990; Reid and Stamps 1997; Stoltz and Andrade 2010; Beckers and Wagner 2011; Robinson et al. 2011; Roff 2015). In these systems, receivers respond to a signal if it is above a threshold value and do not respond if the signal is below the threshold value. For example, female variable field crickets *(Gryllus lineaticeps)* prefer males producing chirps at rates above 3.0 chirps per second but do not discriminate between chirp rate variants that both lie either above or below this threshold (Beckers and Wagner 2011).

Threshold responses may reflect a behavioral decision by receivers (e.g., Beckers and Wagner 2011), but they may also be linked to an animal’s perceptual system. Categorical perception, for example, occurs when continuous variation in a stimulus is perceived as belonging to distinct categories, with individuals showing an increased capacity to discriminate stimuli that fall into different categories as compared to stimuli that fall in the same category (Harnad 1987; Green et al. 2020). For example, a recent study found that female zebra finches *(Taeniopygia guttata)* show categorical perception of the orange to red color continuum representative of male beak color: females labeled eight color variants as lying in two categories and showed better discrimination of cross-category variants as compared to equally-spaced within-category variants (Caves and Green et al. 2018). Categorical perception of signal variation, such as that found in zebra finches (Caves, Green et al. 2018), túngara frogs (Baugh et al. 2008), and sparrows (Nelson and Marler 1989), would likely lead to a threshold response if categories are treated in an binary fashion, thereby violating the continuous assumptions of cost-based models of reliable signaling.

While some models have investigated when different types of threshold responses will evolve (Janetos 1980; Real 1990; Svennungsen et al. 2011; Bleu et al. 2012), few studies have explored how threshold responses influence the evolution of reliable signaling. Many game theoretical approaches to studying reliable signaling have relaxed the assumption that receivers assess continuous variation in signal intensities (Enquist 1985; Maynard Smith 1991; Hurd 1995; Számadó 1999). However, game theoretic models have typically been of discrete signaling games in which both signalers and receivers choose from a discrete set of behaviors (e.g., signalers send either a low- or high-quality signal and receivers either attack or do not attack) or continuous signaling games in both signalers and receivers have a continuum of possible states (Johnstone and Grafen 1992b; Bergstrom and Lachmann 1997). Here, we are interested in the case in which signalers are capable of signaling at any intensity on a continuous range (e.g., tail length can be any length from 5–15mm), but receivers have a binary threshold response (e.g., mate with signaler if and only if tail is longer than 10mm). Broom and Ruxton (2011) have previously explored such a scenario, showing that when receivers respond to signals in a threshold manner it leads to the evolution of “all-or-nothing” signaling systems. That is, signalers with a quality below a critical value all produce the same cheap signal while signalers with a quality above the critical value all produce the same expensive signal. The model of Broom and Ruxton (2011) assumed that receivers all had the same threshold value. However, it is conceivable that receivers could vary in their threshold values, for example due to environmental conditions or developmental history. Here, we explore the evolutionary implications of such inter-individual variation in threshold values.

We begin by developing a model of signal evolution that assumes receivers respond to continuous signal variation in a continuous fashion (akin to the models of Grafen 1990*a*, 1990*b*; Johnstone 1997). This model requires some simplifying assumptions, but it provides a clear demonstration of how reliable signals evolve when there is not a threshold response (the typical assumption). We then show how threshold responses to continuous signal variation can remove the variation in equilibrium signal intensities central to current models of reliable signaling, but how introducing inter-individual variation in threshold responses can rescue the evolution of reliable signals. In addition to these analytical models, we also develop individual-based simulations to assess the robustness of our conclusions under more realistic ecological and genetic assumptions, as well as to ask what role additional complexities in receiver choice can have on the evolution of signals responded to in a threshold fashion.

We focus on the case in which the signaling behavior of the signaler population evolves to optimize fitness and the location of the mean threshold value of the receiver population does not evolve. This assumption is relevant to many biological scenarios in which the threshold value of the receiver population is evolutionarily constrained. For instance, if a receiver’s threshold value is determined by its perceptual machinery rather than a behavioral choice, as with categorical perception, the threshold value might be constrained by its neural physiology (Green et al. 2020; Mason 2020). The threshold value of the receiver population might also be maintained by stabilizing selection from other ecologically important signals and cues. For example, consider a generalist predator (receiver) under selection to avoid eating an aposematically colored toxic prey species (signaler). The threshold value of aposematic coloration above which the predator will not attack an individual of the focal prey species might be constrained by selection to detect the coloration of other prey species. Finally, threshold values might be constrained by a lack of additive genetic variation in the trait or strong genetic linkage with other important phenotypes (Blows and Hoffmann 2005; Gomulkiewicz and Houle 2009). This assumption that the mean threshold value of the receiver population does not evolve simplifies our analytical models, leading to clear predictions. However, we also explore the situation of signaler-receiver coevolution in our individual-based simulations and find that the general conclusions from our analytical models still hold in the presence of signaler-receiver coevolution.

## Continuous Assessment Model

We start by constructing a model of signal evolution that assumes receivers respond to continuous signal variation in a continuous fashion (i.e., there is not a threshold response). This model can be thought of as an intermediate between those of Grafen (1990*a*, 1990*b*) and Johnstone (1997): it can be mathematically difficult to incorporate ecological complexity into Grafen’s models, while Johnstone’s model is purely graphical and thus cannot be analytically evaluated. This model, and our other analytical models below, are rather simple models in which we do not model the evolution of receivers. Instead, we assume that receivers have a fixed reaction norm which is a continuous function that scales positively with signal intensity (i.e., receivers are more likely to respond to signals of greater intensity). We also assume that the optimal signal intensity for a signaler is that which maximizes its net benefit. Therefore, the equilibrium signal intensity is defined as the maximization of the difference between *b*(*s*) and *c*(*s*), where *b*(*s*) is the fitness benefit received by a signaler for producing a signal intensity *s* and *c*(*s*) is the fitness cost associated with producing signal intensity *s*.

In line with previous models (Grafen 1990*a*, 1990*b*; Johnstone 1997), we assume that fitness costs increase linearly with signal intensity and that these costs increase more rapidly for low-quality signalers than high-quality ones (the same qualitative results should apply for any monotonically increasing cost function). Let *c*(*s*) = *α*(*q*)*s*, where *α*(*q*) is a function relating a signaler’s quality, *q,* to its cost of signaling. For analytical tractability, we assume that signaling cost decreases linearly with quality and that 0 ≤ *q* ≤ 1 (*q* = 0 are the lowest quality individuals and *q* = 1 are the highest quality individuals). Therefore,

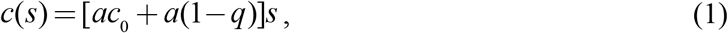

where *a* is the rate at which the slope of the cost function decreases with quality, and *c*_0_ determines the minimum possible cost of signaling, *ac*_0_. Following the graphical representation of Johnstone (1997), we assume that signaler benefit is a saturating function of signal intensity such that

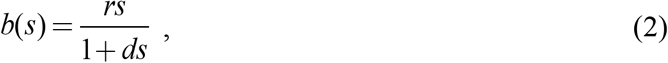

where *r* is the rate that benefit increases with signal intensity and *d* is the degree of saturation with signal intensity (plots of equations [1] and [2] can be seen in Figure 1A). More mechanistically, the benefit function can be thought of as a function of the receivers’ reaction norm, which is not explicitly modeled here but the same qualitative results should apply for any continuous, monotonically increasing benefit function.

**Figure 1.**
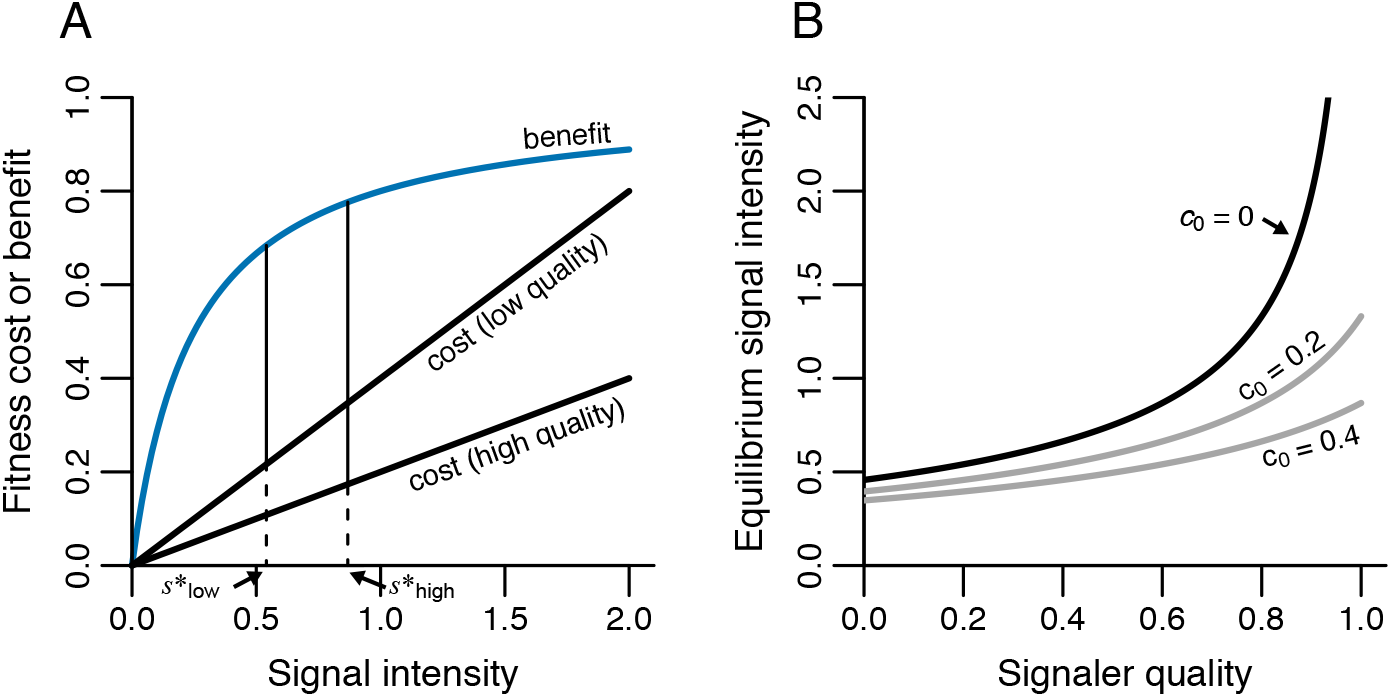
The evolution of reliable signals in a model with continuous assessment. A) Relationship between signal intensity and fitness costs or benefits for signalers of low (*q* = 0.2) and high (*q* = 0.6) quality, given *c*_0_ = 0. Arrows denote the equilibrium signal intensity, *s*,* for low- and high-quality individuals, which occurs where the difference between benefit and cost is the greatest. B) Relationship between signaler quality and equilibrium signal intensity given by equation (3). Three different values for *c*_0_ are shown. Note that higher quality individuals signal at greater signal intensities and that panel A closely matches Johnstone’s graphical model (1997, Figure 7.2). In both panels, *a* = 0.5, *r* = 4, and *d* = 4.

Maximizing the difference between equation (1) and (2) gives the equilibrium signal intensity

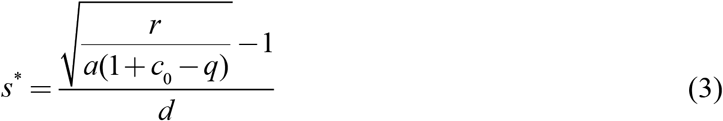

(we provide a derivation in the appendix, sec. A1). This equation shows that higher quality signalers will have greater signal intensities than lower quality ones (Figure 1B). In other words, signal intensity is a reliable indicator of quality. Figure 1A demonstrates this pattern for two different qualities, showing that, for high-quality individuals, the maximum net benefit is at a higher signal intensity than for low-quality individuals. With greater values of *c*, the relationship between signaler quality and signal intensity becomes more linear and less steep (Figure 1B). For simplicity, in all other figures, we assume that *c*_0_ = 0.

The results of this model are in line with those of Grafen (1990*a*, 1990*b*) and Johnstone (1997). This modeling approach also provides a useful framework that can be adapted to analytically explore animal signaling in different biological scenarios, such as threshold assessment.

## Threshold Assessment Model

### Fixed Threshold

We next adapt the above model so that receivers respond to signal variation in a binary fashion (i.e., a threshold response). That is, if a signal is below a threshold value *T*, the receiver assesses the signal as low-quality and if the signal is above *T*, the receiver assesses the signal as high-quality. Because a signaler’s benefit depends on the receiver’s assessment, the benefit of signaling at a given intensity can be thought of as the proportion of the receiver population that assesses the signal as high-quality. For example, in a mating context where males are signaling to females, a male’s benefit of signaling would be the proportion of females in the population that assess it as a high-quality mate and therefore mate with it. Alternatively, in the context of aggressive interactions, the signaler’s benefit is the proportion of competitors that identify the signaler as high-quality and give up a contest against the signaler. We assume that signal intensities at or above the receiver threshold gain maximum benefits, while signal intensities below the threshold gain no benefit. Therefore,

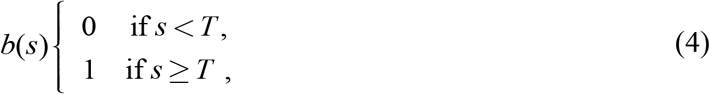

where 1 is the maximum benefit at which every receiver assesses the signaler as high-quality.

In this model, there are only two equilibrium signal intensities: the baseline signal intensity 0 and the threshold value *T.* This is an example of an all-or-nothing signaling system and thus our model agrees with that of Broom and Ruxton (2011). If the costs to an individual of signaling above the threshold are so high that there is never a net benefit of signaling, that individual will signal at the baseline signal intensity of 0. In all other cases, individuals signal exactly at the threshold value (Figure 2A,B). This makes intuitive sense: there is no benefit to signaling below the threshold value because these signals are all assessed as low-quality, but there is also no benefit of signaling any higher than exactly at the threshold value, because this would only accrue additional signaling costs. The result of this effect is that signalers on either side of the threshold will have identical signal intensities of either zero below the threshold or exactly at the threshold value (Figure 2B). In fact, if we assume that for all qualities *q* there is some signal intensity *s* where *b*(*s*) > *c*(*s*), all individuals will signal exactly at the threshold value as is shown in Figure 2C,D. Furthermore, because individuals signaling at an intensity of 0 are never assessed as high-quality, a signal intensity of 0 would only be evolutionarily stable if being assessed as high-quality was not strictly necessary for reproduction. Otherwise, all signalers would signal at the threshold value, irrespective of costs. When all signalers signal at the threshold value (e.g., Figure 2D), there is no information provided by the signal and thus, if receivers’ threshold responses are not ecologically, physiologically, or genetically constrained, we would expect receivers to evolve to ignore the signals all together (an outcome conceptually similar to the lek paradox; Borgia 1979; Tomkins et al. 2004; Kotiaho et al. 2008).

**Figure 2.**
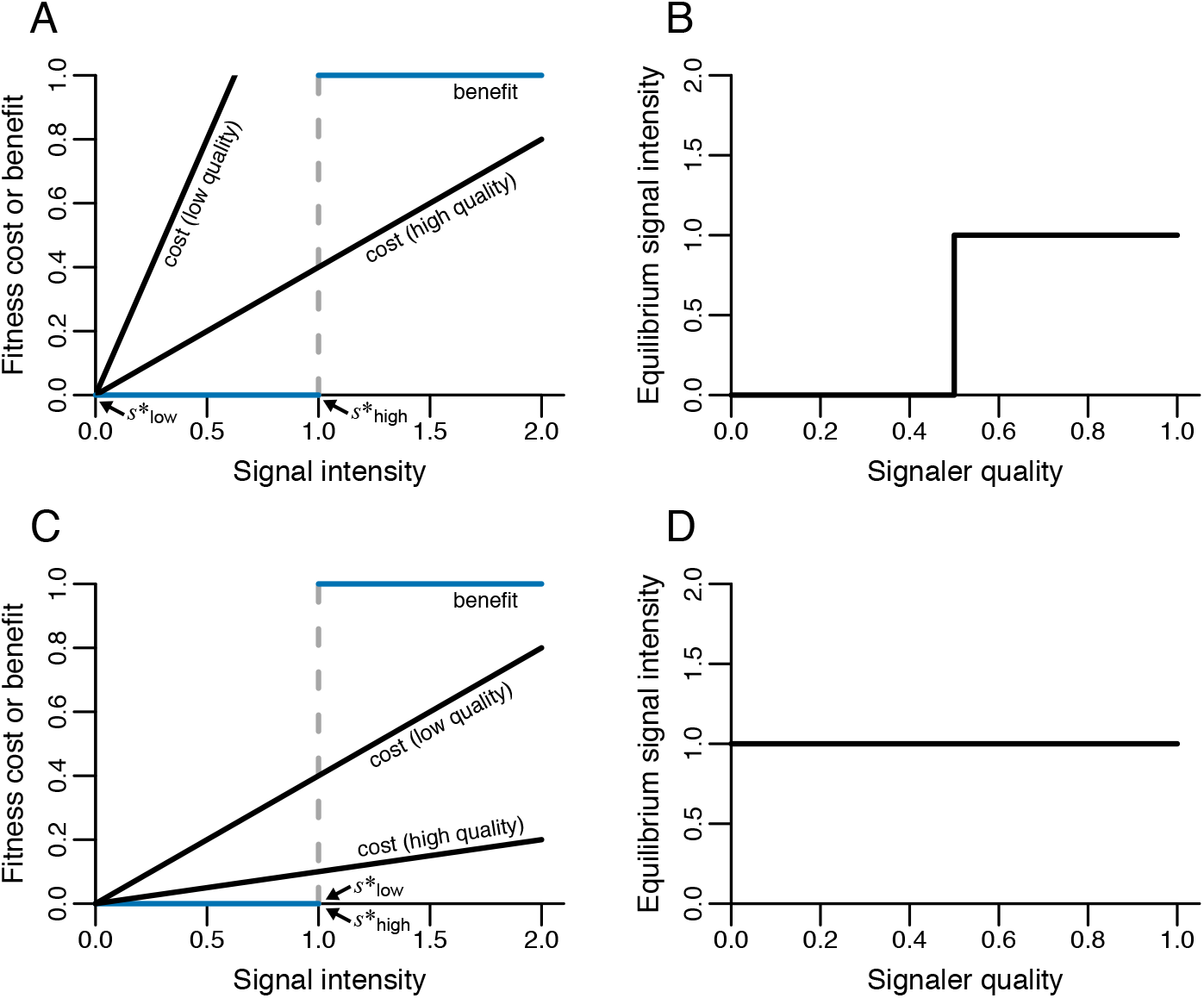
In a model with strict threshold assessment, signalers either evolve ‘all-or-nothing’ signals (A and B) or all signalers signal at the same value (C and D). A and C show graphical depictions of the relationship between signal intensity and fitness costs or benefits for signalers of low (*q* = 0.2) and high (*q* = 0.8) quality. The dashed gray lines indicate the receiver threshold value. Arrows denote the equilibrium signal intensity, *s*,* for low- and high-quality individuals, which occurs where the difference between benefit and cost is the greatest. Note that in C the equilibrium signal intensity is the same for low- and high-quality individuals. B and D show the relationship between signaler quality and signal intensity. In A and B, *a* = 2.0 and in C and D, *a* = 0.5. In all panels, *T* = 1 and *c*_0_ = 0.

We focus on the case of a receiver with a binary response, but it is worth noting that the results of this model can be generalized for categorical responses in which receivers have multiple possible assessment categories (e.g., lowest-quality, low-quality, high-quality, highest-quality). As with the binary case, the number of equilibria signal intensities will be equal to the number of assessment categories.

The results of this model are similar to those of Lachmann and Bergstrom (1998, see also Bergstrom and Lachmann 1997, 1998) who also showed that there are evolutionarily stable signaling strategies in which different quality signalers signal at the same intensity (they refer to these strategies as “pooling equilibria”). However, their approach assumes that only a finite number of signal types are possible, while our model, as well as the model of Broom and Ruxton (2011), allows for a continuum of possible signal intensities and a finite number of signal intensities emerge as a prediction. The models of Lachmann and Bergstrom (1998) and Broom and Ruxton (2011) are in some ways more general than ours because they consider receiver coevolution. By assuming that receiver threshold values are evolutionarily constrained, we were able to obtain similar results with a mathematically simpler model that can now be adapted to consider variation in threshold values.

### Variable threshold

The model above assumes a fixed threshold value for all receivers, but it is likely that threshold values will vary among receivers. For example, the sample of female crickets tested by Beckers & Wagner (2011) showed a threshold of chirp rate for mate choice decisions, but there was also within-sample variance around this threshold value. Similarly, predators may vary in the threshold value of an aposematic signal required to induce avoidance (for examples see Endler and Mappes 2004). Because we assume that receivers do not evolve in this model (which implies threshold values are evolutionarily constrained), our model is most relevant to cases in which variation is threshold values is not genetically determined and rather emerges because of environmental conditions or developmental history (in the *Coevolution of Signalers and Receivers* section below, we evaluate the effects of allowing threshold evolution using individual-based simulations). To model a variable threshold, we now assume that the threshold values of the receiver population are normally distributed with mean 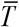 and variance *σ*^2^. Note that this convention implies that receivers can have negative threshold values: if a receiver’s threshold value is less than or equal to 0, it evaluates all signalers as high-quality because the minimum signal intensity is 0. Once again assuming that the benefit of the signal is the proportion of the receiver population that assesses the signal as high quality, it follows that

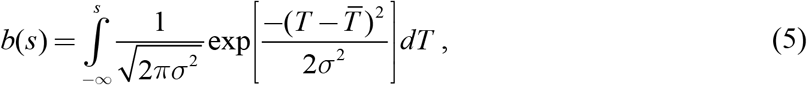

which simply gives that the benefit of signaling at intensity *s* is the area less than *s* under a normal distribution with mean 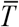 and variance *σ*^2^. For example, if a signal intensity of *s* is assessed as high-quality by 20% of the receiver population, *b*(*s*) = 0.2.

Given equation (5) and assuming that costs still follow equation (1), the equilibrium signal intensity is

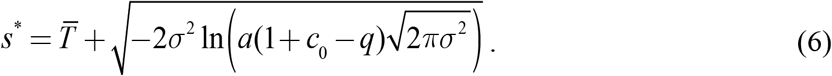

As with equation (3), this equation provides the equilibrium signaling intensity for different qualities of the signalers. However, if *b*(0) > *s*\* or if equation (6) is undefined, which occurs if 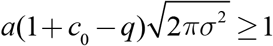, the equilibrium signaling value is instead 0 (see the appendix, sec. A1, for more details and for the derivation of equation [6]).

Equation (6) shows that incorporating variance into the receivers’ threshold values restores the reliability of the signaling system such that signalers of different qualities receive maximum net benefits at different signal intensities (Figure 3A). Furthermore, with greater variance, signal intensity increases faster with increasing quality. That is, for signalers of the same difference in quality, increasing variance in receiver thresholds leads to a larger difference in signal intensity (Figure 3B–D).

**Figure 3.**
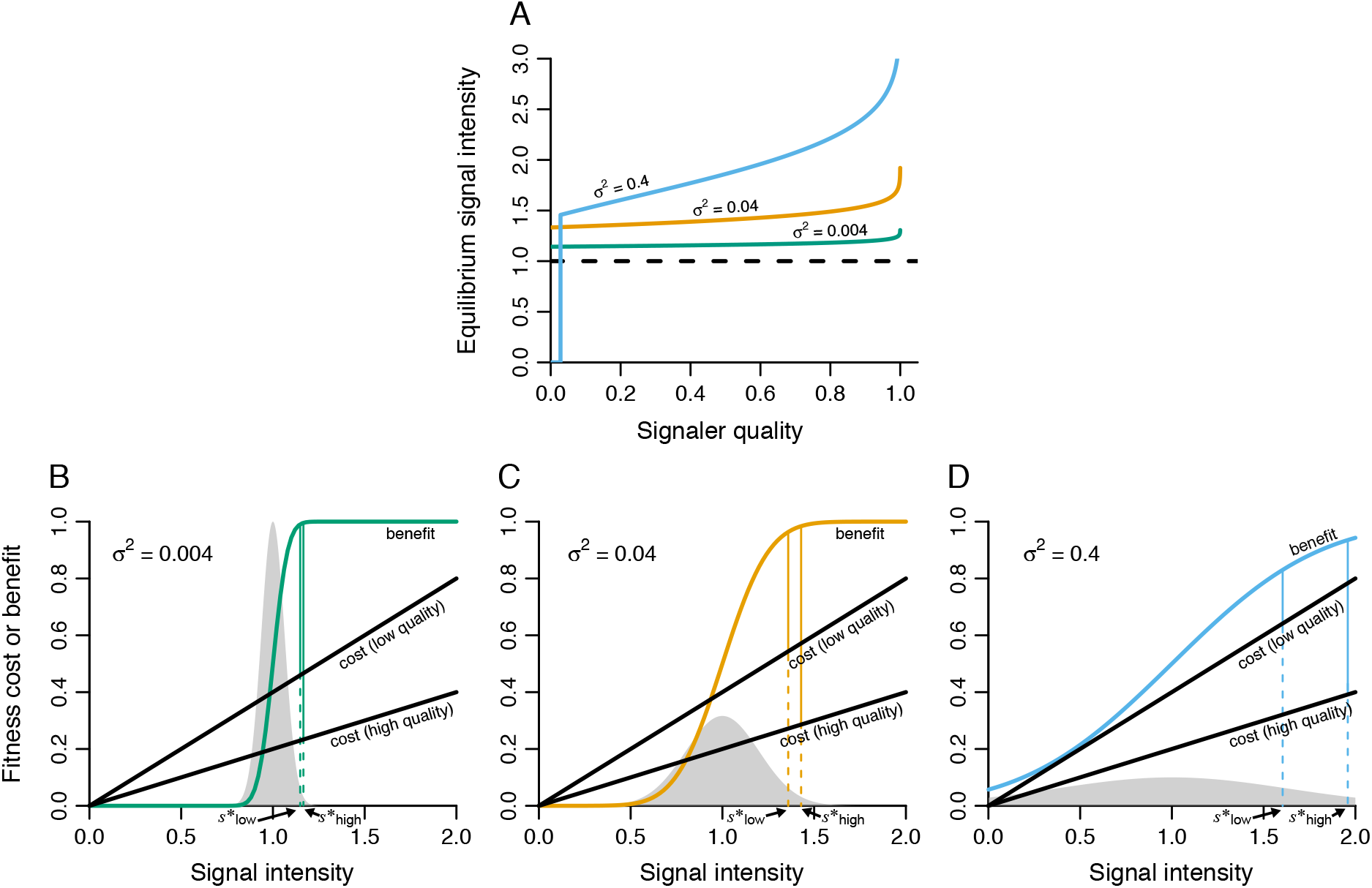
Variation in receiver threshold values restores reliability of signaling systems. A) Relationship between quality and equilibrium signal intensity given by equation (6). Dashed black line denotes the mean receiver threshold value. Colored curves indicate different relationships for different values of variation in threshold value (*σ*^2^). B–D) Relationship between signal intensity and fitness costs or benefits for signalers of low (*q* = 0.2) and high (*q* = 0.6) quality for three different degrees of variation in the threshold value (*σ*^2^). Gray distributions represent the distributions of threshold values in the receiver population. Arrows denote the equilibrium signal intensity, *s*,* for low- and high-quality individuals, which occurs where the difference between benefit and cost is the greatest. Parameters: 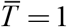, *a* = 0.5, and *c*_0_ = 0.

After introducing inter-individual variation in the threshold value, the only case in which signalers of the different qualities signal at exactly the same value is when they are signaling at the baseline value 0 (e.g., Figure 3A, *σ*^2^ = 0.4 at *q* < 0.028). Because we assume that threshold values follow a normal distribution, there is always at least a small proportion of the receiver population that have a threshold value less than 0 and thus assess all signalers as high-quality. Therefore, there is always some benefit to signaling at the baseline value, and this benefit increases with increased variance in threshold values because more of the receiver population will have a threshold value less than 0.

Interestingly, in this model, the equilibrium signaling value is never between 0 and 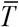 (i.e., equation [6] is always greater than 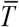 or undefined). In other words, individuals either evolve to signal at the baseline value of 0 or to signal at some intensity above the mean threshold value 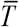. This is surprising because, in this model, there is a benefit to signaling between 0 and the mean threshold value. However, further exploration reveals that the net benefit is always greater to signal either above 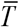 or at 0. This is due to the sigmoidal shape of the benefit function: below 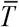, the benefit function curves towards lower values, but above 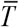 it curves towards higher values (Figure 3B–D). The sigmoidal shape of the benefit function emerges as a result of the normal distribution of the receiver variance. Because costs are linear, this sigmoidal shape results in the net benefit being greater at values above 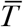 than below it. If the variation in receiver thresholds has a different distribution, however, it is possible for an equilibrium signaling value to fall between 0 and 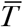 (see the appendix [sec. A3, fig. A1], for an example assuming a gamma distribution).

## Individual-based Simulations

The analytical models above make many simplifying assumptions (e.g., ignoring genetics and assuming infinite population sizes). To assess the robustness of our results and to explore further ecological complexity, we developed stochastic individual-based simulations. These simulations track individuals’ genotypes and phenotypes and model the evolution of signaling behavior. Unlike in the analytical models, these simulations actually model adaptation via changes in allelic values rather than simply maximizing the net benefit. For these simulations we chose to model the evolution of a mating signal in a sexually reproducing species with two sexes (signalers and receivers). In principle, this same modeling framework could be adapted to fit other types of signaling systems (e.g., intraspecific competition or predator-prey interactions).

### Methods

Each run of the simulation modeled a population in which each individual was either a signaler or a receiver. Each signaler had a quality *q* (ranging from 0 to 1), which determined its cost of signaling. Each receiver had a threshold *T*, which was used to evaluate signalers (details below). Generations were overlapping and population size was regulated by the number of mating sites available *K.* The simulation was broken up into time steps, which we will refer to as years. There was an annual sequence of events which happened in the following order: mortality, filling of mating sites, reproduction (including signal assessment and mating, Figure 4).

**Figure 4.**
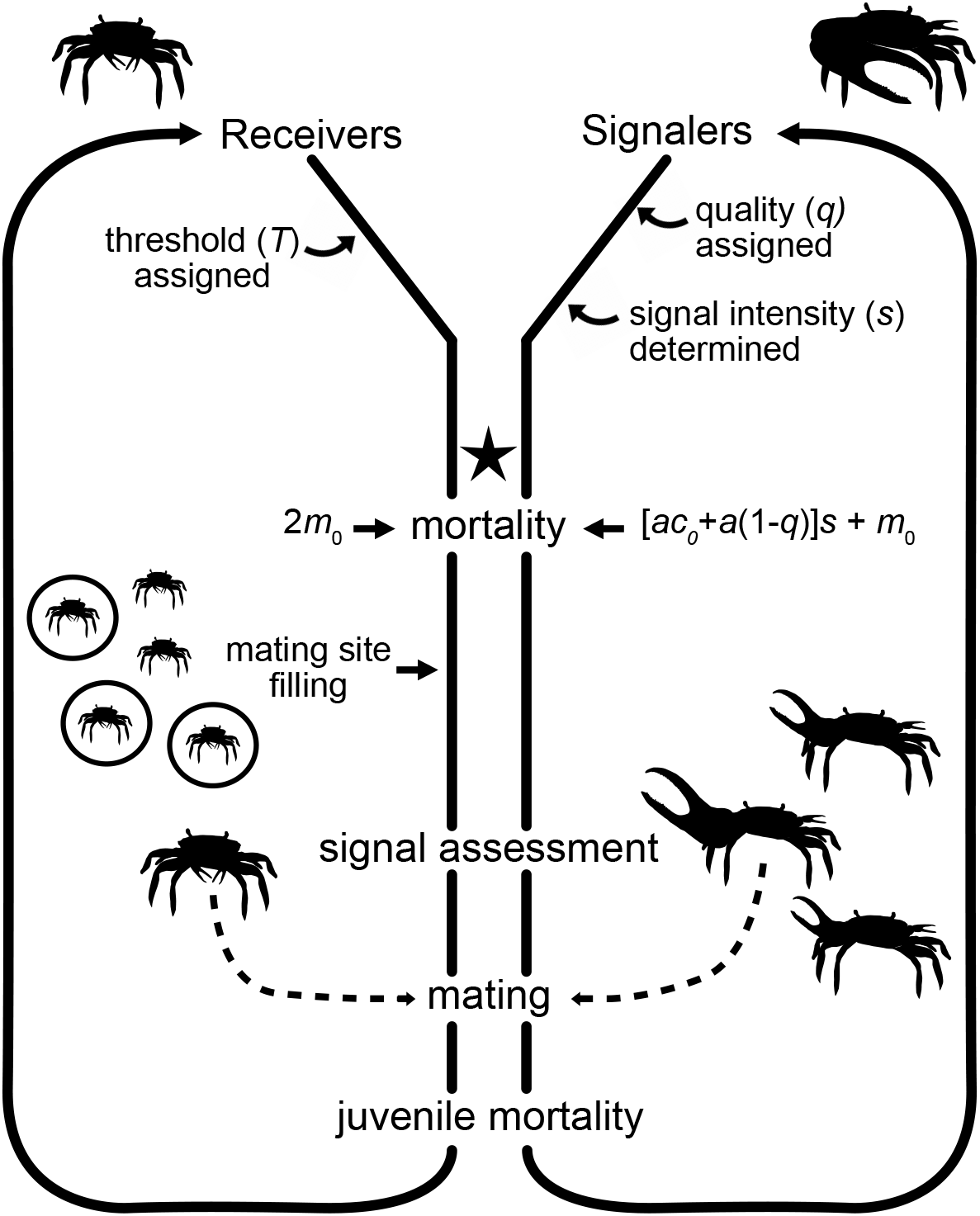
Life-history diagram for the individual-based simulations. The star indicates the start of the annual cycle. Silhouettes of fiddler crabs (genus *Uca)* are included as visual examples for signalers and receivers in a mating context.

Mortality occurred at the start of each year independent of age or reproductive status. For signalers, quality and signal intensity affected mortality such that *M* = [*ac*_0_ + *a*(1 – *q*)]*s* + *m*, where *m* is the baseline mortality, *a* is the rate at which the slope of the cost function decreases with quality, *c*_0_ determines the minimum cost of signaling, and *s* is the signal intensity. Receivers died with a fixed mortality of 2*m*_0_, which was higher than the baseline signaler mortality in order to keep sex ratios relatively balanced.

Receivers occupied mating sites, from which they evaluated the signals of signalers. There were only *K* mating sites available each year. If there were more than *K* receivers in the population, *K* receivers were randomly selected (without replacement) to occupy mating sites that year. Unselected receivers did not mate that year. We assumed that there was no spatial structure to the population such that all individuals had the same probability of arriving at any site and all sites became unoccupied at the end of the year.

Receivers selected signalers as mates by evaluating their signal intensity, *s*. Receivers had threshold assessment of signals such that if a signaler’s signal intensity was above the receiver’s threshold value *T*, the receiver evaluated the signaler as a high-quality mate and if the signal intensity was below *T*, the receiver evaluated the signaler as low-quality. Receivers mated with the first signaler they evaluated as high-quality and did not mate again that year. Each year, receivers evaluated up to *N* signalers. If by that point no signalers were evaluated as high-quality, the receiver mated with the final signaler that it evaluated regardless of quality (i.e., threshold with last-chance option; Janetos 1980). The threshold value of each receiver was randomly assigned at birth (regardless of genotype) from a normal distribution with mean 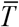 and variance *σ*^2^. Each mated pair produced *B* offspring per year and all offspring survived with a probability 0.5.

Each signaler had a trait, *z*(*q*), which determined its signal intensity as a function of its quality. This trait was genetically determined by 11 additive diploid loci, each with infinitely many possible allelic values. Each locus determined the signaling value when the signaler had a specific quality (broken up by increments of 0.1). That is, locus 1 determined the signal intensity when the signaler’s quality was 0.0, locus 2 determined the signal intensity when the signaler’s quality was 0.1, and so on. This implementation allows signal intensity to be almost any function of quality. The phenotypic values of signal intensity for a given quality were determined by adding together the genotypic values of each haplotype at the respective locus plus a random environmental component drawn from a zero-mean normal distribution with variance *e*^2^. If an individual’s genotypic value was less than 0, its phenotype was assigned to be 0. An individual’s phenotype (signal intensity) was determined at birth and did not change throughout its lifetime. Note that, because *q* is a continuous quantity, but the trait *z*(*q*) is in increments of 0.1, signalers evaluated their quality by rounding to the nearest increment of 0.1. Therefore, signalers could not perfectly evaluate their own quality, which is a realistic assumption for many organisms (e.g., Percival and Moore 2010).

At birth, each offspring was randomly assigned to either the signaling sex (signalers) or the receiving sex (receivers) and each signaler was randomly (uniform distribution, 0–1) assigned a quality *q*. Each haplotype of the offspring was assigned by randomly selecting one of the alleles at each locus of the respective parent (i.e., independent recombination). During birth, a mutation occurred on each locus with probability *μ*. If a mutation occurred, a random value drawn from a zero-mean normal distribution with variance *ρ*^2^ was added to the value at that locus. Receivers were only carriers for this trait, and it did not affect their preference.

Each run of the simulation was initiated with 2*K* individuals whose genotypes were randomly assigned from a normal distribution with mean 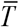 and variance 0.2 (sex, quality, and signal were randomly assigned in the same manner as they were assigned at birth). To allow populations to reach selection-mutation-drift balance, there was a 10,000-year “burn-in” period at the beginning of the simulation. We used two different methods for this burn-in period. In the first, receivers randomly selected mates independent of signal and there was no cost of signaling. This simply allowed the population to reach mutation-drift balance so that the initially assigned genotypes did not affect results. In the second method, receivers assessed signals in a continuous manner, where the probability of mating with a signaler was [(*rs*) / (1 + *ds*)] + 0.05, where 0.05 is the baseline probability of mating and *r* and *d* are the same as in equation (2). This latter method models how signals evolve if receivers evolve threshold assessment after previously having had continuous assessment. In both cases, following the 10,000-year burn-in period, simulations then ran for 20,000 years with threshold assessment. Results were qualitatively similar using both methods for the 10,000-year burn-in period. For simplicity, all presented results are for the case in which receivers responded in a continuous fashion during the burn-in period because there was less variation among runs of those simulations (see figure A2 for results of simulations with random mating during the burn-in period).

### Results

The results of our individual-based simulations matched the qualitative predictions of our analytical model. As predicted by the analytical model, with little variance in threshold values, signalers evolved to signal at relatively similar values whether they were low- or high-quality; however, with greater variance, the difference between signal values became greater (Figure 5).

**Figure 5.**
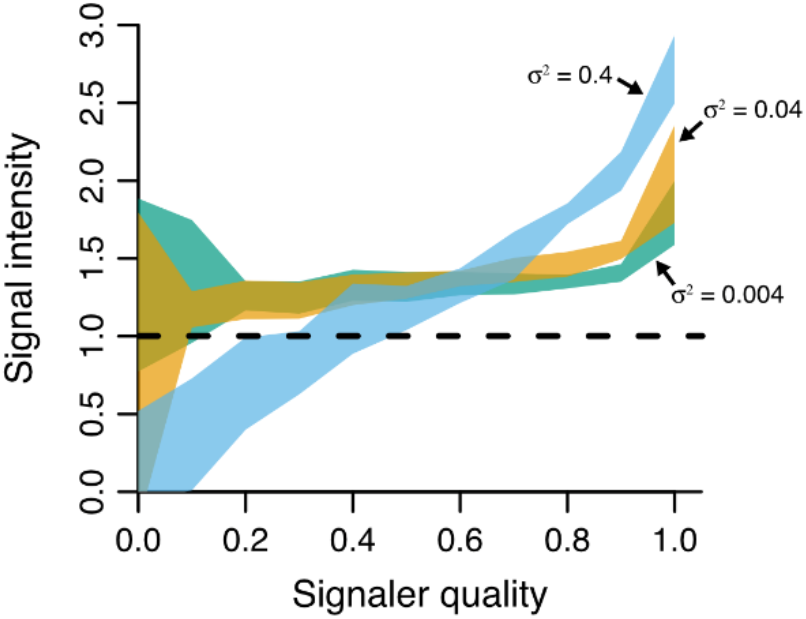
The relationship between signaler quality and signal intensity in the individual-based simulations. Compare to the analytical model results shown in Figure 3A. The dashed black line denotes the mean threshold value. Shaded areas represent the mean of 10 runs of the simulation plus or minus one standard deviation. Different colored shading represents different levels of threshold variation (*σ*^2^). Negative genotypic values were interpreted as zeros, as that is how they affected phenotype. Parameters: 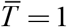, *a* = 0.5, *c*_0_ = 0, *K* = 300, *u* = 0.25, *ρ*^2^ = 0.01, *e*^2^ = 0.0001, *B* = 5, *N* = 10.

There was greater variance in the signal intensity of low-quality individuals compared with high-quality ones both within and among runs of the simulation (among run variation shown in Figure 5). This is likely because there was more genetic drift at the loci associated with signaling when low-quality. Because low-quality individuals were less likely to mate, there were fewer opportunities for selection to act on these loci. Although low-quality individuals signaled at lower intensity than high-quality ones on average, in some runs of the simulation, genetic drift led to low-quality individuals signaling at unexpectedly high signal intensities.

The individual-based simulations provide additional information about the phenotypic distribution of signalers in the population that cannot be gained by our analytical methods (Figure 6). These distributions show that even if the mean genotype of the population was to always signal above the threshold value (as was the case for *σ*^2^ = 0.004 and *σ*^2^ = 0.04 in Figure 5), there were still individuals in the populations with signal intensities below the threshold value. In other words, as long as there is variation in threshold values, signal intensities can vary along a continuum (Figure 6). Furthermore, the greater the variation in threshold values, the more uniform the distribution of signal intensities (Figure 6). For all simulation parameters, there was a peak in the number of individuals signaling at an intensity of 0. This is because 0 is the minimum signal intensity and thus the tail of the phenotypic distribution that would be below 0 was all clustered into that phenotypic value.

**Figure 6.**
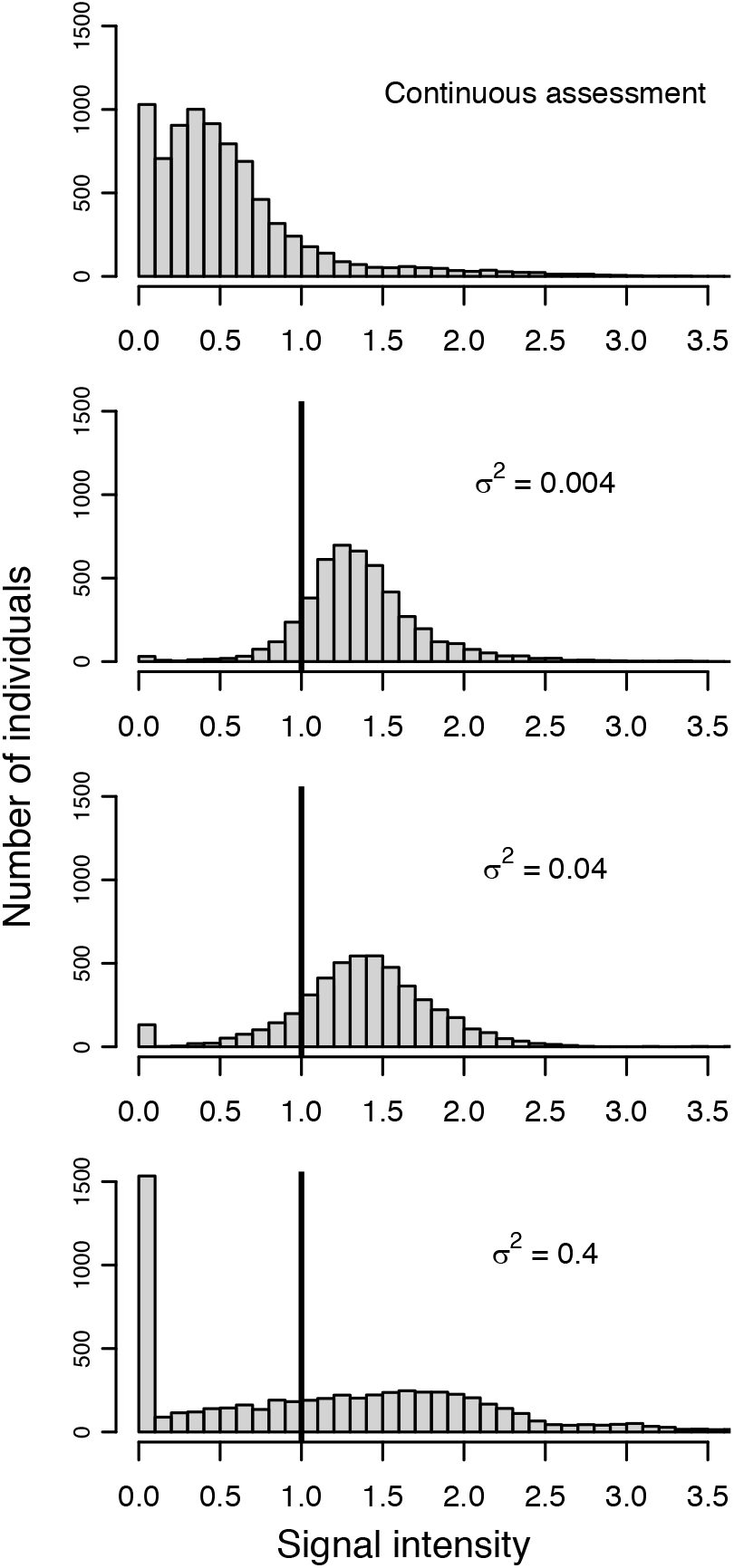
The phenotypic distribution of signalers in the individual-based simulations. Each panel shows the distributions for the cumulative number of individuals in 5 runs of the simulation. The solid vertical line indicates the mean threshold value of the receiver population. Parameters: 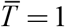, *K* = 300, *u* = 0.25, *ρ*^2^ = 0.01, *e*^2^ = 0.0001, *B* = 5, *N* = 10.

One major difference between our analytical model and individual-based simulations is that the simulations did not show the sudden change from signaling at the baseline value to signaling above the threshold value that was seen the analytical models (compare *σ*^2^ = 0.4 in Figure 3A and Figure 5). This disagreement can be explained by considering the ecological assumptions of our models. Recall in the analytical model that there was always some benefit to signaling at the baseline value because threshold values were normally distributed and thus some proportion of the receiver population assessed all signal intensities as high-quality. In essence, this logic assumes that the population is infinitely large, which was obviously not the case in the individual-based simulations.

However, in our individual-based simulations, a benefit of signaling at the baseline value emerged in another way, which was by receivers mating with the final signaler that they evaluated regardless of its quality (i.e., the last-chance option). This provided signalers some chance of mating, even if their signal intensity was below the threshold value of all receivers. The lower the maximum number of signalers a receiver evaluated *N*, the greater this baseline benefit, because there was an increased chance that any given signaler was the final one a receiver evaluated. Reducing the value of *N* in the simulations resulted in sudden changes in signal intensities similar to those seen in the analytical model (Figure 7). Furthermore, the lower the value of *N*, the higher the quality at which this sudden change occurred. This pattern is intuitive because with a greater baseline benefit, populations should evolve to only signal above the baseline value when there is a larger net-benefit, which occurs at high qualities. Indeed, in the extreme case of *N* = 1, which would amount to random mating, we would expect all individuals to signal at the baseline value and thus sustain the minimum signaling costs.

**Figure 7.**
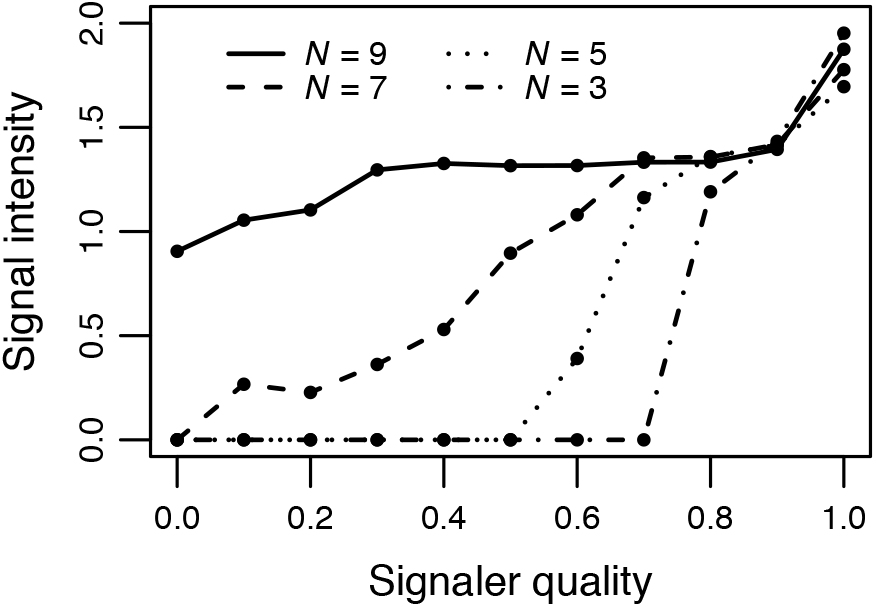
Results of the individual-based simulations for different values for the maximum number of mates a receiver could evaluate *N*. If a receiver did not encounter a signaler with a signal intensity above the receiver’s threshold, it mated with the final signaler it evaluated regardless of signal intensity. Each line shows the mean genotypic values for 10 runs of the simulation. Parameters: 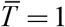, *σ*^2^ = 0.004, *K* = 300, *u* = 0.25, *ρ*^2^ = 0.01, *e*^2^ = 0.0001, *B* = 5.

We also restructured simulations so that receivers did not mate with the last mate that they evaluated and instead did not mate at all if none of the *N* signalers evaluated had a signal intensity above their threshold value (i.e., fixed-threshold without a last-chance option; Roff 2015). In these simulations, mean signaling values below the threshold value were less likely to evolve (Figure A3). This is logical, because as with increasing the value of *N* in the simulations with a last-chance option, this scenario decreases the benefit of signaling at the baseline value. The only way a signaler can mate is if its signal intensity is above the threshold value of a receiver.

We also ran simulations in which variation was incorporated into the signalers instead of the receivers. Biologically, this occurs when a signal (a phenotype) is influenced by local environment conditions (i.e., phenotypic plasticity). In these simulations, all receivers had the same threshold value and we incorporated variation into the signalers by increasing the random environmental component of the phenotype *e*^2^, which is equivalent to decreasing the heritability of the signaling trait. As with variation in threshold values, in these simulations, signals once again evolved so that higher quality individuals signaled at greater signal intensities than lower quality ones, and the strength of this pattern increased with more variation in the random environmental component (Figure A4). This result occurs because there is selection for high-quality individuals to signal farther above the threshold value to ensure that their phenotype (and their offspring’s phenotypes) are above the threshold value regardless of the environmental effect. As above, in these simulations, receivers mated with the last signaler evaluated, so there was some benefit for low-quality individuals to signal below the threshold value because they could reduce the costs of signaling and still have a chance of being the last signaler a receiver evaluated.

### Coevolution of Signalers and Receivers

Up to this point, all of our models have assumed that the mean threshold value of the receiver population does not evolve. In this section, we present results from a model that we adapted from our individual-based simulations to explore the coevolution of signalers and receivers. To do so, we assumed that a receiver’s threshold value was determined by a single-locus quantitative trait and that receivers had greater fecundity if they mated with a higher quality signaler. These simulations were not intended to investigate all of the nuances of signaler-receiver coevolution, but instead were intended to test whether the general conclusions from our above models hold even with receiver evolution. These simulations were identical to the individual-based simulation described above except for the key differences mentioned below.

The receiver’s threshold value was determined by a single-locus quantitative trait. All individuals (signalers and receivers) had a genotypic value for this trait, but it was only expressed in receivers. There was no genetic correlation between threshold genotypes and signaling genotypes. Each receiver’s phenotypic value of the threshold was their genotypic value plus a random environmental component drawn from a zero-mean normal distribution with variance *σ*^2^.

Recall that in the original model, receivers did not acquire any fitness benefits for mating with higher quality signalers. In this version of the model, however, we wanted threshold values to evolve, so we included fitness benefits for mating with higher quality signalers. This was implemented by having receivers have one additional offspring (*B* + 1 offspring) with a probability of their mate’s quality (recall that quality ranges from 0 to 1).

The genotype (for the threshold trait) of offspring was simply the average value of both of their parents. To speed up the pace of evolution, we assumed that a mutation occurred with every birth such that a random value drawn from a zero-mean normal distribution with variance 0.02 was added to the offspring’s genotype. These mutations added additional variance to the threshold values, a factor that was not included in our previous models (the analytical model or other simulations). This difference precludes quantitative comparisons of our previous models with the coevolutionary models we present here, but we are still able to make qualitative comparisons.

Simulations were initiated as in previously described simulations. Unless otherwise specified (see figure A5 for exceptions), individuals’ initial threshold values were randomly assigned with a mean of 1 and variance 0.04. For the 10,000-year “burn-in” period, individuals had continuous assessment after which they evaluated signalers in a threshold manner. After the burn-in period, simulations ran for 20,000 generations. This simulation length appeared to be long enough for thresholds to reach a stable value (Figure A5).

As in our previous models, these simulations showed that increased variance in threshold values leads to a greater difference between the signaling values of low- and high-quality signalers (Figure A6), thus demonstrating that the qualitative results of our previous models are unchanged, even if receivers’ thresholds and signalers’ signals are allowed to coevolve.

## Discussion

Most previous models of signal evolution have assumed that receivers assess continuous variation in a signals in a continuous manner (e.g., Grafen 1990*a*, 1990*b*; Johnstone 1994). In many species, however, receivers exhibit threshold responses in which receivers respond to continuously varying signals in an binary fashion (e.g., Masataka 1983; Zuk et al. 1990; Reid and Stamps 1997; Stoltz and Andrade 2010; Beckers and Wagner 2011; Robinson et al. 2011). We have shown that (1) invariant threshold assessment of signals leads to all-or-nothing signals (Broom and Ruxton 2011) or a breakdown of reliable signaling systems, but (2) reliable signaling can be restored if variation is introduced either in the value of the threshold boundary among receivers or in the translation of genotype to phenotype among signalers. In addition, (3) when reliable signaling evolves, the mean signal intensity of signalers will typically be above the mean threshold value of receivers, but (4) the population of signalers will still show a continuum of signal intensities from well below to well above the population mean threshold.

Our models emphasize the importance of variation among receivers in maintaining reliable signaling systems. In natural populations, it seems highly likely that threshold values will vary among receivers to some degree because local environmental conditions affect phenotypes. Indeed, in a mating context, there is often considerable variation in mate choice among receivers due to genetic, developmental, or environmental differences (Jennions and Petrie 1997). In animals exhibiting categorical perception in particular, there is evidence of variation in category boundaries (Caves and Green et al. 2018; Zipple and Green et al. 2019; Caves et at. 2020) as well as context-dependence of categorical boundaries (Lachlan and Nowicki 2015). Future empirical work should quantify the degree of variation among receivers’ threshold values and more directly investigate its effect on signal evolution. Empirical work could also test conclusions (3) and (4) above by comparing the distribution of signal intensities and threshold values in threshold signaling systems.

Although our models explicitly considered signals whose reliability is maintained as a result of high- and low-quality signalers facing differential costs of signaling, a similar framework also applies to signals whose reliability is maintained as a result of high- and low-need signalers receiving differential benefits of signaling (Johnstone 1997). An example of this latter case would be nestling birds begging for food from their parents. In this scenario, the benefit of signaling (begging) increases more rapidly for high-need nestlings than low-need ones. As is the case for models with continuous assessment, the same general conclusions emerge from our threshold assessment models whether we assume signalers have differential costs or benefits (see the appendix [sec. A2, fig. A7 and fig. A8] for differential benefits results).

Similarly, while we have only discussed among-receiver variation in threshold values (i.e., inter-receiver variation), the same general patterns should hold if individual receivers vary in their threshold values over time (intra-receiver variation). This intra-receiver variation might emerge from changes during development or changes in environmental conditions, but it could also be a result of variation in an individual receiver’s ability to detect a threshold boundary. That is, a receiver might mistakenly assess a signal as above its threshold when in actuality it was not or *vice versa.* Our analytical model treats all these types of variation identically; therefore, it predicts that either inter- or intra-receiver variation in threshold values will maintain signal reliability. This suggests that while perceptual errors by the receiver decrease the number of equilibrium signaling values in models with continuous assessment (Johnstone and Grafen 1992a), in models with threshold assessment, perceptual errors might in fact *increase* the number of equilibrium signaling values. Our individual-based simulations only explicitly modeled inter-receiver variation, however. It would be useful for future studies to explore intra-receiver variation more explicitly.

Our analytical models focus on pure fixed threshold assessment, in which individual receivers evaluate any signal above their threshold value as high-quality and below their threshold as low-quality. However, there are different variants of threshold assessment in which receivers responses follow a general threshold rule but are context-dependent (reviewed by Roff 2015). For instance, there is evidence that female variable field crickets, *Gryllus lineaticeps,* evaluate mates using a threshold strategy with a last-chance option in which females use a threshold rule, but if a female has evaluated *N* males and none of them were above the threshold value, it mates with the final male encountered (Janetos 1980; Beckers and Wagner 2011). The results of our individual-based simulations show that while our general conclusions about signal evolution hold whether receivers follow as a pure fixed threshold strategy or have a last-chance option, the specific relationship between signaler quality and signal intensity will differ depending on the strategy. Moreover, with the last-chance option, this relationship depends on the number of signalers evaluated before mating with the final signaler. We have not evaluated other context-dependent threshold strategies (reviewed in Roff 2015), but it is likely that these will also alter the specific relationship between signaler quality and signal intensity.

In addition, we have only considered the evolution of signals under relatively simple ecological scenarios in which reliable signaling is maintained by differential signaling costs (or benefits), but in more complicated scenarios, reliable signals can evolve even without immediate signaling costs. For example, reliable signaling can evolve without signaling costs when there are repeated interactions (Rich and Zollman 2016). Similarly, reliable signaling might evolve via kin selection when individuals are signaling among close relatives (e.g., begging in birds; Caro et al. 2016). Furthermore, our models do not explain the evolution of dishonest signals (in our models, signals either evolve to be reliable or to contain no information). Dishonest signals (or “bluffs”) can be maintained in otherwise reliable signaling systems by the existence of frequency dependent selection (Számadó 2017), a factor that is not present in our models.

Another complexity we do not consider here is the ability of receivers to assess their own quality. Mutual assessment plays and important role in signaling, particularly in the context of animal contests where individuals might have to compare their own resource holding potential to that advertised by their opponent (Arnott and Elwood 2009; Elwood and Arnott 2012). When evaluating the signals of opponents, an individual might have a threshold response, such that its giving-up decision is based on a threshold signal value set by the individual’s own resource holding potential. Understanding how threshold responses affect signal evolution in such situations would be a valuable future direction of our work. It is worth noting, however, that despite many examples of mutual displays before contests, there is surprisingly little empirical evidence that contestants compare their own display to the display of their opponent (Elwood and Arnott 2012).

It is important to recognize that most of our models assume there is a fixed mean threshold value for the receiver population. If variations in threshold values among receivers are due to heritable differences, threshold values themselves will be subject to natural selection and potentially change over time (Janetos 1980; Real 1990; Bleu et al. 2012). We would expect the mean threshold value to remain stable if threshold values are either physiologically constrained, as might be the case in some species with categorical perception, or ecologically constrained, as might be the case when the same perceptual machinery is used to assess signals in addition to being used in some other ecological context (e.g., female guppies, *Poecilia reticulata,* use the color orange for mate assessment and food detection; Rodd et al. 2002).

The coevolution of threshold and signal does not change the qualitative results of our models. Our individual-based simulations with coevolution confirmed that the qualitative conclusions of our models can still hold when selection causes receivers’ thresholds to evolve (Figure A6). However, we only explored a limited set of ecological and genetic assumptions in those simulations, and additional work is necessary to more fully understand how the coevolution between signalers’ signals and receivers’ thresholds affects the reliability of signaling systems. For instance, in intra-specific signaling systems (e.g., mate choice), there is often a genetic correlation between receiver preference (here, the threshold value) and signaling traits, which can lead to emergent evolutionary phenomena such as Fisherian runaway selection (Fisher 1930; Lande 1981; Kirkpatrick 1982). Future studies should explore how threshold responses by receivers affects these processes.

## Author Contributions

All authors conceptualized the study together and contributed to model design and interpretation. J.H.P. constructed the models and wrote the first draft of the manuscript. P.A.G., M.N.Z., and S.N. contributed to the writing of the manuscript.

## Acknowledgments

We are grateful to E.M. Caves, S. Peters, S. Johnsen, and S. Zlotnik for providing value advice throughout this project. We also thank N. Kortessis and the members of the UF SRNE journal club for providing comments on earlier versions of this manuscript. Funding was provided by the Duke University Office of the Provost. PAG was supported by a Human Frontier Science Program Fellowship # LT000460/2019-L. M.N.Z. was supported by an NSF graduate research fellowship.

## Data Accessibility Statement

Code for the individual-based simulations along with accompanying documentation will be posted on Dyrad upon acceptance. The code is available to reviewers upon request.

