## appendix for "Threshold assessment, categorical perception, and the evolution of reliable signaling"

### A1. Derivations of equilibrium signal intensities

*Continuous assessment model (equation [3])*

As is stated in the main text, our analytical models assume that the equilibrium signaling intensity is the signal intensity with the greatest net benefit. Let $f(s)$ be the net benefit of a signal intensity of $s$, such that

$$f(s) = b(s) - c(s) \quad \text{(A1)}$$

$$f(s) = \frac{rs}{1+ds} - [a(1+c_0-q)]s \quad \text{(A2)}$$

The maximum value of this function is where $f'(s) = 0$ and $f''(s) < 0$. These derivatives are

$$f'(s) = \frac{r}{(1+ds)^2} - [a(1+c_0-q)] \quad \text{(A3)}$$

$$f''(s) = \frac{-2dr}{(1+ds)^3}. \quad \text{(A4)}$$

Setting equation A3 equal to zero and solving for $s$ gives

$$s = \frac{-1 \pm \sqrt{\dfrac{r}{[a(1+c_0-q)]}}}{d}. \quad \text{(A5)}$$

Note that, as long as $d$, $r$, and $s$ are all positive, equation (A4) is always negative. Thus, equation (A5) always provides the maximum net benefit. Furthermore, because the term in the square root is always positive (assuming $0 < q < 1$) and negative values of $s$ are not biologically reasonable, equation (A5) can be simplified to equation (3) in the main text, which gives the signal intensity that obtains the maximum net benefit.

*Threshold Assessment model (equation [6])*

As above, let $f(s)$ be the net benefit of a signal intensity of $s$, which in this model is given by

$$f(s)=\int_{-\infty}^{s}\frac{1}{\sqrt{2\pi\sigma^2}}\exp\left[\frac{-(T-\bar{T})^2}{2\sigma^2}\right]dT-[a(1+c_0-q)]s\,. \tag{A6}$$

To calculate the derivatives, we must assume that assume that the lower bound of the integral is a constant. Therefore, we assume that $T$ > -1,000,000. If $\bar{T}$ is positive and variance is not extremely large, as is assumed throughout this manuscript, this truncation will have a negligible effect on the statistical properties of the distribution. Given this assumption, the first and second derivatives are respectively,

$$f'(s)=\frac{1}{\sqrt{2\pi\sigma^2}}\exp\left[\frac{-(s-\bar{T})^2}{2\sigma^2}\right]-[a(1+c_0-q)] \tag{A7}$$

$$f''(s)=\frac{-(s-\bar{T})\exp\left[\frac{-(s-\bar{T})^2}{2\sigma^2}\right]}{\sigma^3\sqrt{2\pi}}\,. \tag{A8}$$

Setting equation (A7) equal to zero and solving for $s$ gives

$$s=\bar{T}\pm\sqrt{-2\sigma^2\ln\left(a(1+c_0-q)\sqrt{2\pi\sigma^2}\right)} \tag{A9}$$

Recall, equation (A9) is only a maximum when $f''(s)<0$. For evaluation of the sign of $f''(s)$, we can ignore the denominator because it will always be positive. In addition, the exponential term will always be positive. Therefore, $f''(s)<0$ if and only if

$$\bar{T}<s \tag{A10}$$

From this, we can simplify equation (A9) to equation (6) in the main text (when $\bar{T} > s$, [A9] gives the minimum net benefit, which is not of interest for our questions).

However, unlike equation (3), equation (6) does not always give the signal intensity with the maximum net benefit. This is because the maximum can also be at the signal intensity of 0, in which case $f'(s) \neq 0$. Therefore, to find the maximum, one must compare *b*(0) to equation (6) and take the signal intensity with the higher value (note that *b*(0) = *f*(0) because *c*(0) always equals 0) . Examples of when equation (6) does and does not predict the optimal phenotype are shown in figure A9. Also note that equation (A9) (and thus equation [6]) will be undefined when $a(1+c-q)\sqrt{2\pi\sigma^2} \geq 1$. In these scenarios, the net benefit will always be a decreasing function of signal intensity, and thus the maximum signal intensity will be 0.

**A2. Differences in benefits (need) instead of costs**

Similar results emerge from our models if instead of assuming that different quality individuals face different signaling costs, we assume that different need signalers receive different benefits of signaling. In these models, we assume that quality does not affect cost and that $c_0 = 0$, therefore costs are described by $c(s) = as$.

*Continuous assessment model*

For this model, we assume that benefit depends on need following the equation

$$b(s) = \frac{\nu rs}{1+ds}\ , \qquad \text{(A11)}$$

where $\nu$ is an individual's need. Solving for the maximum net benefit as was done in part 1 of the appendix gives that the optimal signal intensity is

$$s^* = \frac{\sqrt{\frac{\nu r}{a}} - 1}{d} \quad . \tag{A12}$$

From this, we see that reliable signal evolves in the model with differential benefits, such that higher need individuals signal at a greater signal intensity (Figure A7).

*Threshold assessment model*

With a fixed (invariant) threshold, the models with different costs and benefits yield identical results, such that there are only two equilibrium signal intensities: the baseline signal intensity 0 and the threshold value *T*.

In this model with inter-individual variation in the threshold value (with the same assumptions as in the variable threshold section of the main text), benefit is given by the function

$$b(s) = \left( \int_{-\infty}^{s} \frac{1}{\sqrt{2\pi\sigma^2}} \exp\left[ \frac{-(T-\overline{T})^2}{2\sigma^2} \right] dT \right) \nu \, . \tag{A13}$$

Solving for the maximum net benefit and simplifying as was done in part 1 of the appendix gives that the optimal signal intensity is

$$s^* = \overline{T} + \sqrt{-2\sigma^2 \ln\left( \frac{a\sqrt{2\pi\sigma^2}}{\nu} \right)} \quad . \tag{A14}$$

This shows that reliable signaling evolves in a threshold model with a variable threshold and different needs (Figure A8). As with equation (A9), equation (A14) does not always provide the optimal signal intensity because it can also be at the value of 0. Therefore, one must compare $b(0)$ to equation (A14) and take the signal intensity with the higher value as the optimal signal intensity. Also note that equation (A14) will be undefined when $\left( a\sqrt{2\pi\sigma^2} \right) / \nu \geq 1$.

**A3. Threshold assessment model with gamma distributed threshold values**

This model is the same as the analytical variable threshold assessment model in the main text, but we change the assumption that threshold values of the receiver population are normally distributed and instead assume that they follow a gamma distribution. Biological traits are often gamma distributed, so this a realistic assumption. With this assumption, the benefit for signaling at intensity $s$ is given by the function

$$b(s) = \int_0^s \frac{\beta^k T^{k-1} e^{-\beta T}}{(k-1)!} dT \ , \qquad \text{(A15)}$$

where $k$ and $\beta$ are the shape and rate parameters of a gamma distribution, respectively. The net benefit $f(s)$ is thus

$$f(s) = \int_0^s \frac{\beta^k T^{k-1} e^{-\beta T}}{(k-1)!} dT - [a(1 + c_0 - q)]s \, , \qquad \text{(A16)}$$

where the second term is the cost function given by equation (1). The first and second derivatives of $f(s)$ are, respectively,

$$f'(s) = \frac{\beta^k s^{k-1} e^{-\beta s}}{(k-1)!} - [a(1 + c_0 - q)] \qquad \text{(A17)}$$

$$f''(s) = \frac{\beta^k s^{k-2} e^{-\beta s}(k - \beta s - 1)}{(k-1)!} \ . \qquad \text{(A18)}$$

Setting equation (A17) equal to zero and solving for $s$ gives

$$s = \frac{-(k-1)W\left(\frac{-\beta\left(\sqrt[k-1]{(k-1)!\beta^{-k}[a(1+c_0-q)]}\right)}{k-1}\right)}{\beta}, \tag{A19}$$

where $W(z)$ is the Lambert $W$ function ($W(z)$ has two solutions when $-1/e \leq z < 0$, in this range, our equation refers to the lower value, i.e., the lower branch of the function). Equation (A19) gives the maximum of equation (A16) and thus the equilibrium signaling value when equation (A18) is negative, which occurs when $k - \beta s - 1 < 0$.

The mean of a gamma distribution is $k / \beta$ and the variance is $k / \beta^2$. Figure A1 provides plots of equation (A19) for different values of $k$ and $\beta$. These show that our general conclusions from the main text hold and that for some parameter values (low $k$ and $\beta$, which gives high variance relative to the mean) the equilibrium signaling value can be between 0 and the mean threshold value. However, the shape of the distributions of threshold values that lead to equilibrium signaling intensities between 0 and the mean threshold value (e.g., Figure A1D) do not seem as biologically likely as the ones that lead to equilibrium signaling intensities always being 0 or above the threshold value (e.g., Figures 3A and Figure A1B–C).

Equation (A19) is rather complicated, so we note that for large $k$ the gamma distribution converges to a normal distribution with mean $k / \beta$ and variance $k / \beta^2$. Therefore, for large values of $k$, the equilibrium signal intensity is well approximated by the much simpler equation (6). We also note that when $k = 1$ the gamma distribution is an exponential distribution with mean $1/\beta$ and variance $1/\beta^2$. For an exponential distribution, the equilibrium signaling value is given by the more manageable function

$$s^* = \frac{-\ln\left[\frac{[a(1-c_0-q)}{\lambda}\right]}{\lambda} \quad , \tag{A20}$$

(derivation not shown) where $\lambda$ is the rate parameter of an exponential distribution. For an exponential distribution, the second derivative of the net benefit function is $\lambda^2(-e^{-\lambda s})$, which is always negative, and thus equation (A20) always gives the maximum net benefit in this scenario.

### Supplementary Figures

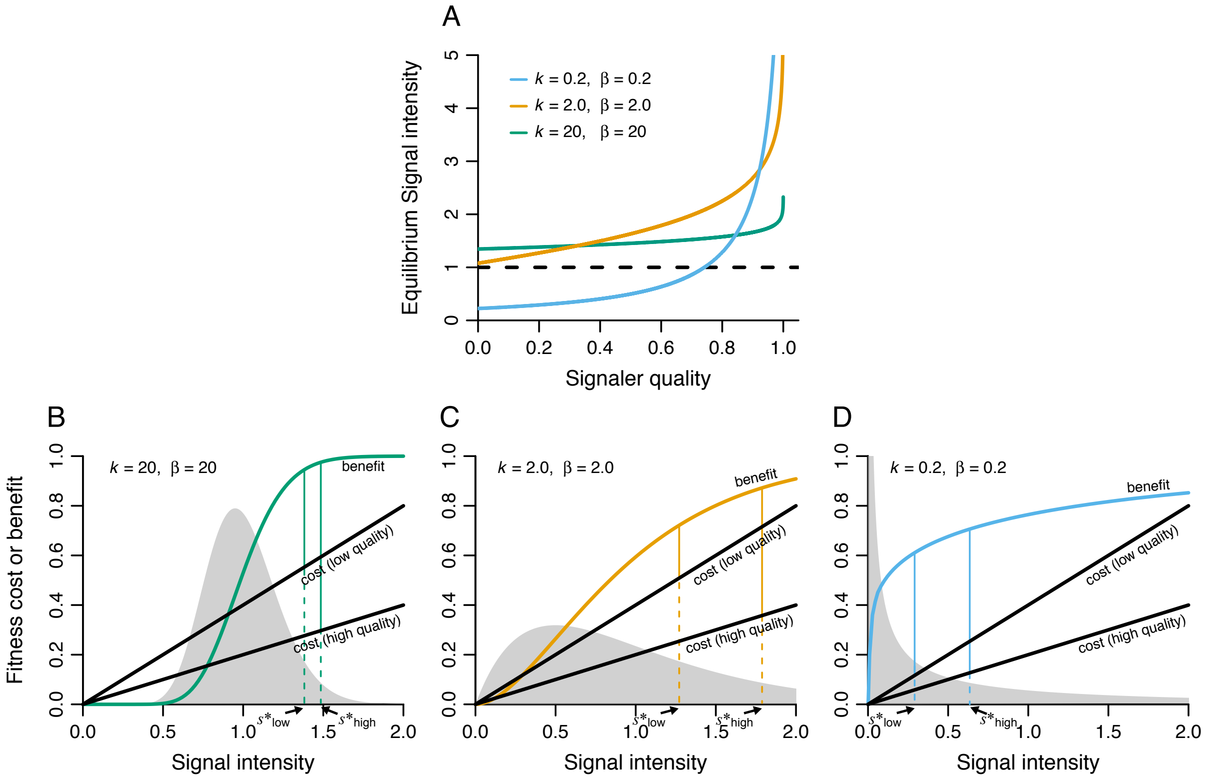


**Figure A1.** Evolution of signaling systems when threshold values of the receiver population follow a gamma distribution. A) Relationship between quality and equilibrium signal intensity given by equation (A19). Colored curves indicate different values for the shape $k$ and rate $\beta$ parameters of a gamma distribution. Dashed black line denotes the mean threshold value ($k / \beta$), which was the same for all parameter combinations shown. B–C) Relationship between signal intensity and fitness costs or benefits for signalers of low ($q$ = 0.2) and high ($q$ = 0.6) quality for the three different parameter sets shown in in panel A. Gray distributions represent the distributions of threshold values in the receiver population (recall variance of a gamma distribution is $k / \beta^2$). Arrows denote the equilibrium signal intensity, $s^*$, for low- and high-quality individuals, which occurs where the difference between benefit and cost is the greatest. Parameters: $a = 0.5$, and $c_0 = 0$.

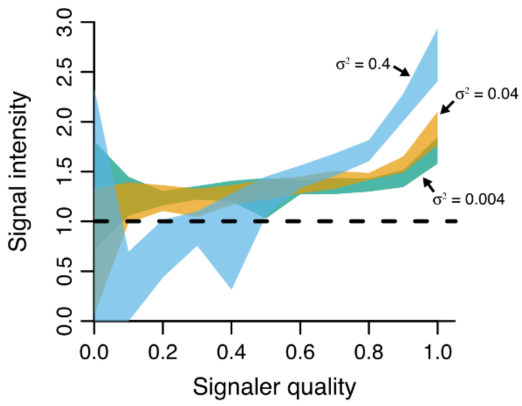


**Figure A2.** The relationship between quality and signal intensity in the individual-based simulations with random mating and no cost of signaling during the 10,000-year burn-in period. Dashed black line denotes the mean threshold value. Shaded areas represent the mean of 10 runs of the simulation plus or minus one standard deviation. Negative genotypic values were interpreted as zeros, as that is how they affected phenotype. Note that results are nearly identical to those in Figure 5. Parameters: $\overline{T} = 1$, $a = 0.5$, $c_0 = 0$, $K = 300$, $u = 0.25$, $\rho^2 = 0.01$, $e^2 = 0.0001$, $B = 5$, $N = 10$.

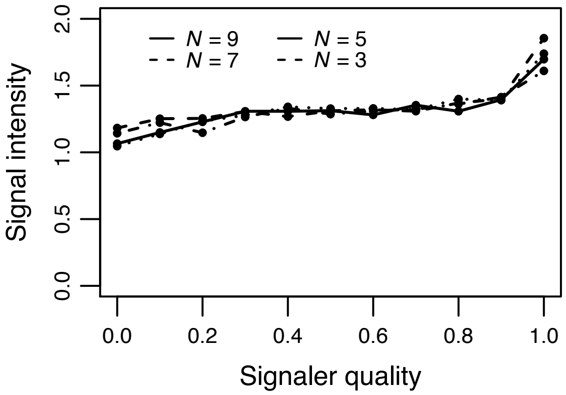


**Figure A3.** Results of individual-based simulations with no last-chance option. Each line shows results for different values for the maximum number of mates a receiver could evaluate *N*. Compare results with Figure 7. In these simulations, many populations went extinct. Each line shows the mean genotypic values for 10 runs of the simulation in which the populations persisted for the entire run of the simulation. Parameters: $\overline{T} = 1$, $\sigma^2 = 0.004$, $K = 300$, $u = 0.25$, $\rho^2 = 0.01$, $e^2 = 0.0001$, $B = 5$.

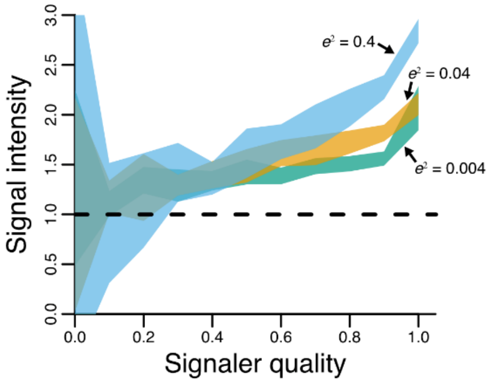


**Figure A4.** Results of the individual-based simulations for different degrees of variance in the environmental component of the phenotype (signal intensity) $e^2$. Shaded areas represent the mean of 10 runs of the simulation plus or minus one standard deviation. Dashed black line denotes the threshold value. Negative genotypic values were interpreted as zeros, as that is how they affected phenotype. Parameters: $\sigma^2 = 0$, $K = 300$, $u = 0.25$, $\rho^2 = 0.01$, $B = 5$, $N = 10$.

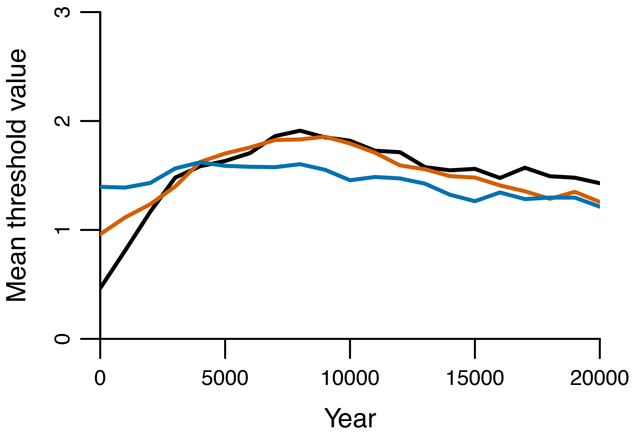


**Figure A5.** Evolution in mean threshold values in the individual-based simulations with coevolution of signalers' signals and receivers' thresholds for three different initial mean threshold values: 0.5 (black), 1.0 (orange), 1.5 (blue). Each line shows the mean of 10 runs of the simulation. Parameters: $K = 300$, $u = 0.25$, $\rho^2 = 0.01$, $e^2 = 0.0001$, $B = 5$, $N = 10$, $\sigma^2 = 0.004$.

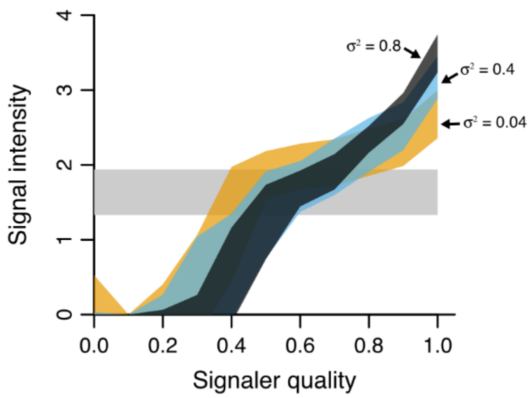


**Figure A6.** Results of the individual-based simulations with coevolution between signalers' signals and receivers' thresholds. Shaded areas represent the mean of 20 runs of the simulation plus or minus one standard deviation. Colors (orange, blue, black) indicate different degrees of variation in the optimal phenotype ($\sigma^2$), here implemented as variance in the environmental component of the threshold phenotype. Negative genotypic values were interpreted as zeros, as that is how they affected phenotype. Shaded gray area indicates the mean threshold value of all 60 simulations plus or minus one standard deviation. The threshold values for all simulations are shown together because the mean values for different degrees of variation were all similar. Parameters: $K = 300$, $u = 0.25$, $\rho^2 = 0.01$, $B = 5$, $e^2 = 0.0001$, $N = 10$.

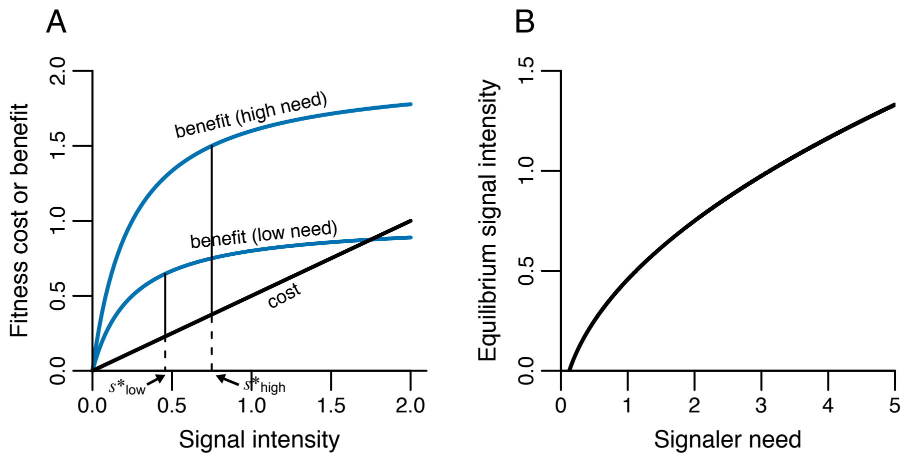


**Figure A7.** The evolution of reliable signals in a model with different needs and continuous assessment. A) Relationship between signal intensity and fitness costs or benefits for signalers of low ($\nu = 1$) and high ($\nu = 2$) need. Arrows denote the equilibrium signal intensity, $s^*$, for high- and low-quality individuals, which occurs where the difference between benefit and cost is the greatest. B) Relationship between signaler need and signal intensity given by equation (A12). Note that higher need individuals signal at greater signal intensities and that panel A closely matches Johnstone's (1997) graphical model. In both panels, parameters were $a = 0.5$, $r = 4$, and $d = 4$.

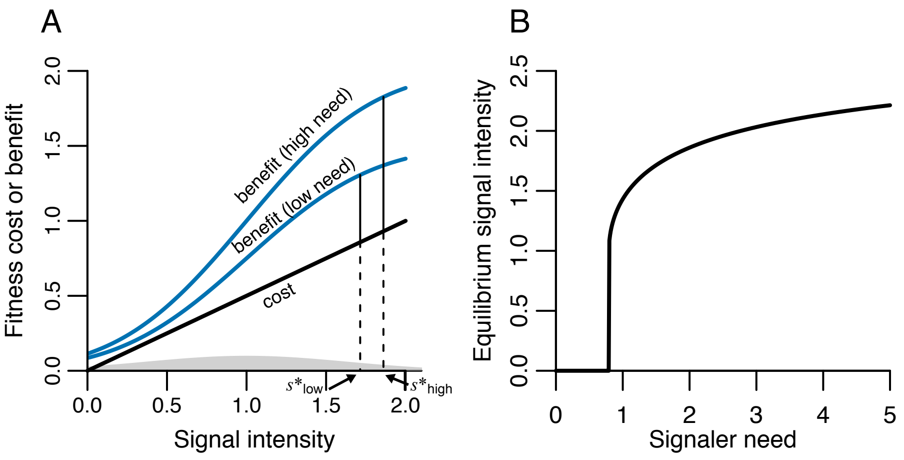


**Figure A8.** The evolution of reliable signals in a model with different needs and threshold assessment with inter-individual variation in signaler's threshold values. A) Relationship between signal intensity and fitness costs or benefits for signalers of low ($\nu = 1.5$) and high ($\nu = 2$) need. Arrows denote the equilibrium signal intensity, $s^*$, for high- and low-quality individuals, which occurs where the difference between benefit and cost is the greatest. B) Relationship between signaler need and signal intensity given by equation (A14). Note that higher need individuals signal at greater signal intensities. In both panels, parameters were $\overline{T} = 1$, $a = 0.5$, $\sigma^2 = 0.4$.

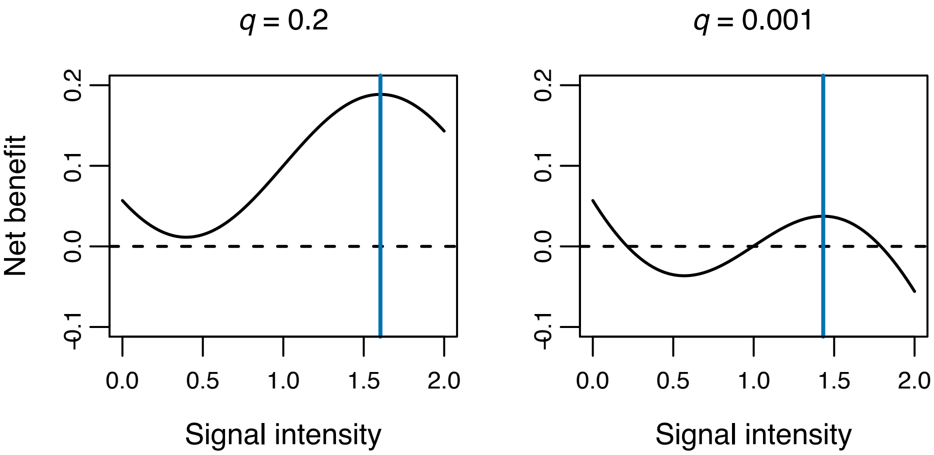


**Figure A9.** Examples of net benefit as a function of signal intensity as given by equation (A6). Vertical blue lines indicate the optimal signal intensity given by equation (6) in main text. The left panel is an example in which equation (6) correctly predicts the optimal signal intensity. The right panel is an example in which equation (6) does not correctly predict the optimal signal intensity because the highest net benefit occurs at a signal intensity of 0, at which $f'(s) \neq 0$. In both panels parameters were $\overline{T} = 1$, $\sigma^2 = 0.4$, $a = 0.5$, and $c_0 = 0$.
